# Genome-wide developed microsatellites reveal a weak population differentiation in the hoverfly *Eupeodes corollae* (Diptera: Syrphidae) across China

**DOI:** 10.1101/606657

**Authors:** Meng-Jia Liu, Xiao-qiang Wang, Ling Ma, Li-Jun Cao, Hong-Ling Liu, De-Qiang Pu, Shu-Jun Wei

## Abstract

The hoverfly, *Eupeodes corollae,* is a worldwide natural enemy of aphids and a plant pollinator. To provide insights into the biology of this species, we examined its population genetic structure by obtaining 1.15-GB random genomic sequences using next-generation sequencing and developing genome-wide microsatellite markers. A total of 79,138 microsatellite loci were initially isolated from the genomic sequences; after strict selection and further testing of 40 primer pairs in eight individuals, 24 polymorphic microsatellites with high amplification rates were developed. These microsatellites were used to examine the population genetic structure of 96 individuals from four field populations collected across southern to northern China. The number of alleles per locus ranged from 5 to 13 with an average of 8.75; the observed and expected heterozygosity varied from 0.235 to 0.768 and from 0.333 to 0.785, respectively. Population genetic structure analysis showed weak genetic differentiation among the four geographical populations of *E. corollae*, suggesting a high rate of gene flow reflecting likely widespread migration of *E. corollae* in China.

## Introduction

*Eupeodes corollae* is one of the most common hoverflies with a worldwide distribution [1, 2]. The larval stage of this species is mostly insectivorous, feeding mainly on aphids [3-5] while adults are pollinators [6-8]. Many hoverfly species are important biological control agents of aphids due to their rapid dispersal and absence of summer diapause compared with other aphidophaga [9]. Understanding the biology and behavior of hoverflies can help in assessing their potential as biological control agents of aphids.

Hoverflies migrate seasonally as revealed by radar monitoring [10] and isotopic tools [11]. Population genetic analysis is also frequently employed to reveal the migration of species as a complementary approach to traditional methods [12-15]. In populations of the hoverflies *Cheilosia longula* [16], *Blera fallax* [17], *Sphaerophoria scripta* and *Episyrphus balteatus* [18], population genetic differentiation has not been found between some regions, suggesting migratory movements of these hoverflies between regions including southern and northern regions of Europe [18, 19]. However, some hoverflies, such as *E. balteatus* and *Scaeva selenitica*, are only partially migratory [20].

Previous studies reported that *E. corollae* is a highly migratory species in Europe [21-23], but its migratory behavior of *E. corollae* remains unclear in other areas. *E. corollae* is commonly found across China, but the ecology and biology of this species has rarely been studied [8]. In this study, we conducted a preliminary examination of the population genetic structure of *E. corollae* in China. First, we obtained random genomic sequences of *E. corollae* using next-generation sequencing and developed an effective and informative set of microsatellite markers of *E. corollae*. We used this novel set of microsatellite markers to investigate the genetic structure of four *E. corollae* populations collected from four representative regions across China.

## Materials and methods

### Sample collection and DNA extraction

A male adult from a laboratory (Sichuan Academy of Agriculture Sciences)-reared line of *E. corollae* was used for generating genome sequences. Four field populations of *E. corollae* were collected from China in March to July 2017 (Table 1, Figure 1a). To avoid the sampling of siblings, adults in a site were collected using insect net with individuals sampled separated by about 20 meters. A total of eight individuals from field collections were used for initial testing of selected primers. Twenty-four individuals from each of the four populations were then used for a population level survey. All samples were stored in absolute ethanol, frozen at −80 °C and stored at the Integrated Pest Management Laboratory of the Beijing Academy of Agriculture and Forestry Sciences. The thorax from each individual *E. corollae* was used for genomic DNA extraction using DNeasy Blood & Tissue Kit (Qiagen, Hilden, Germany).

**Table 1.**
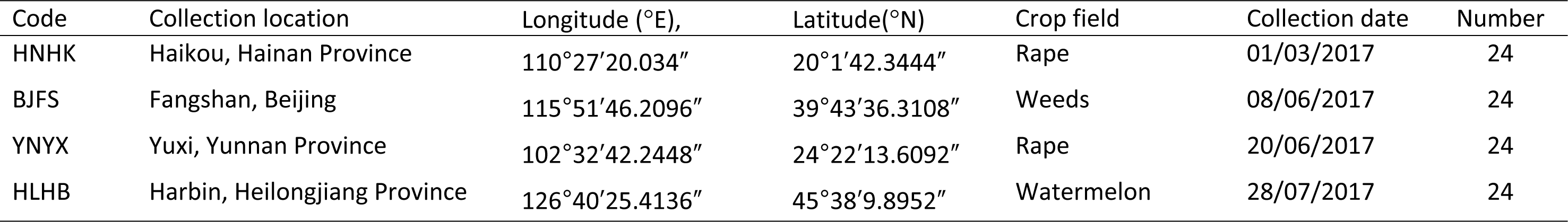
Collection information of *Eupeodes corollae* for microsatellite development and population genetic structure analysis.

| Code | Collection location | Longitude (°E), | Latitude(°N) | Crop field | Collection date | Number |
| --- | --- | --- | --- | --- | --- | --- |
| HNHK | Haikou, Hainan Province | 110°27'20.034" | 20°1'42.3444" | Rape | 01/03/2017 | 24 |
| BJFS | Fangshan, Beijing | 115°51'46.2096" | 39°43'36.3108" | Weeds | 08/06/2017 | 24 |
| YNYX | Yuxi, Yunnan Province | 102°32'42.2448" | 24°22'13.6092" | Rape | 20/06/2017 | 24 |
| HLHB | Harbin, Heilongjiang Province | 126°40'25.4136" | 45°38'9.8952" | Watermelon | 28/07/2017 | 24 |

**Table 2.**
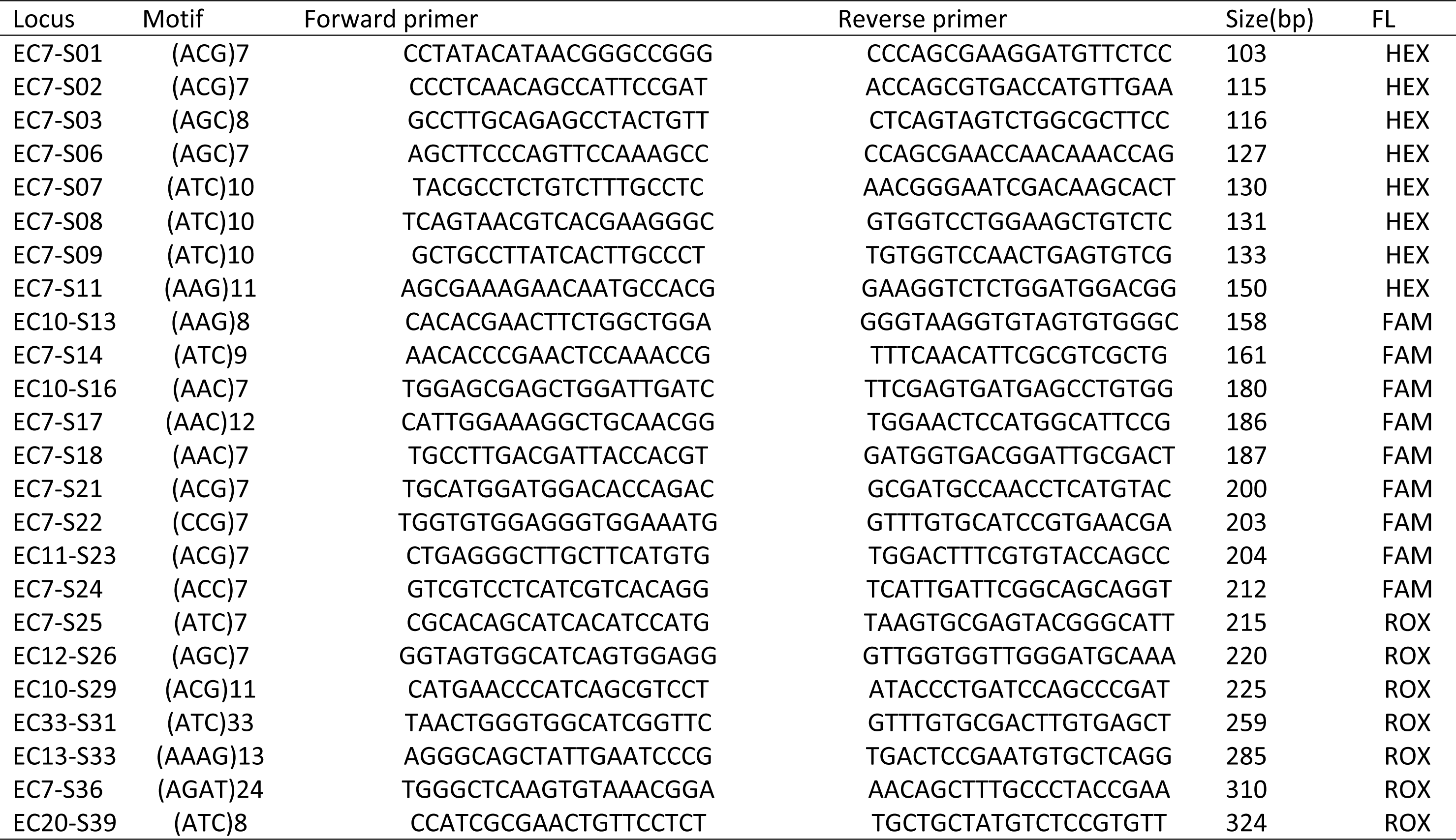
Twenty-four microsatellite loci developed for *Eupeodes corollae*.

**Table 3.**
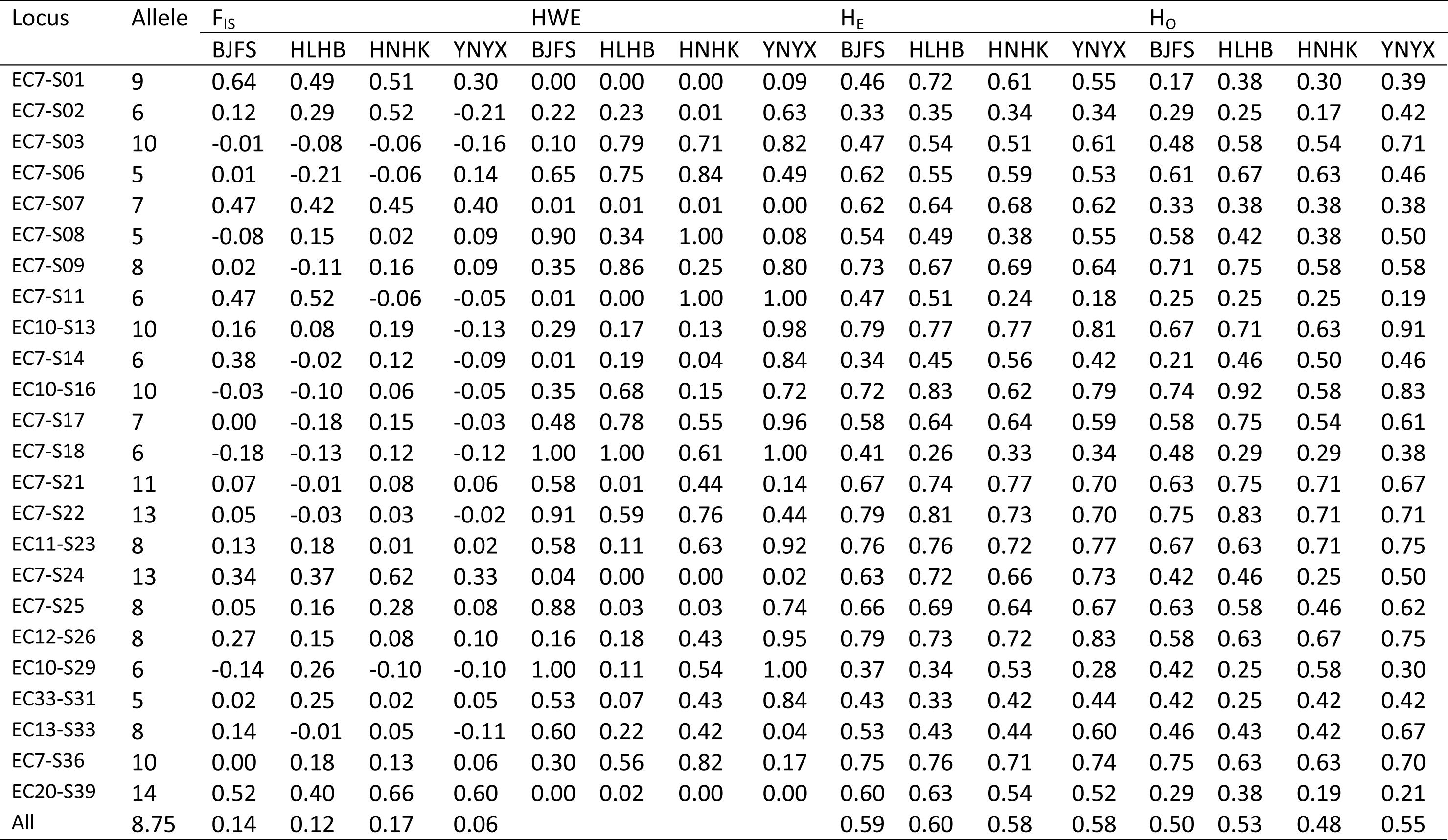
Summary statistics of 24 microsatellite markers for *Eupeodes corollae* validated in four populations.

**Figure 1.**
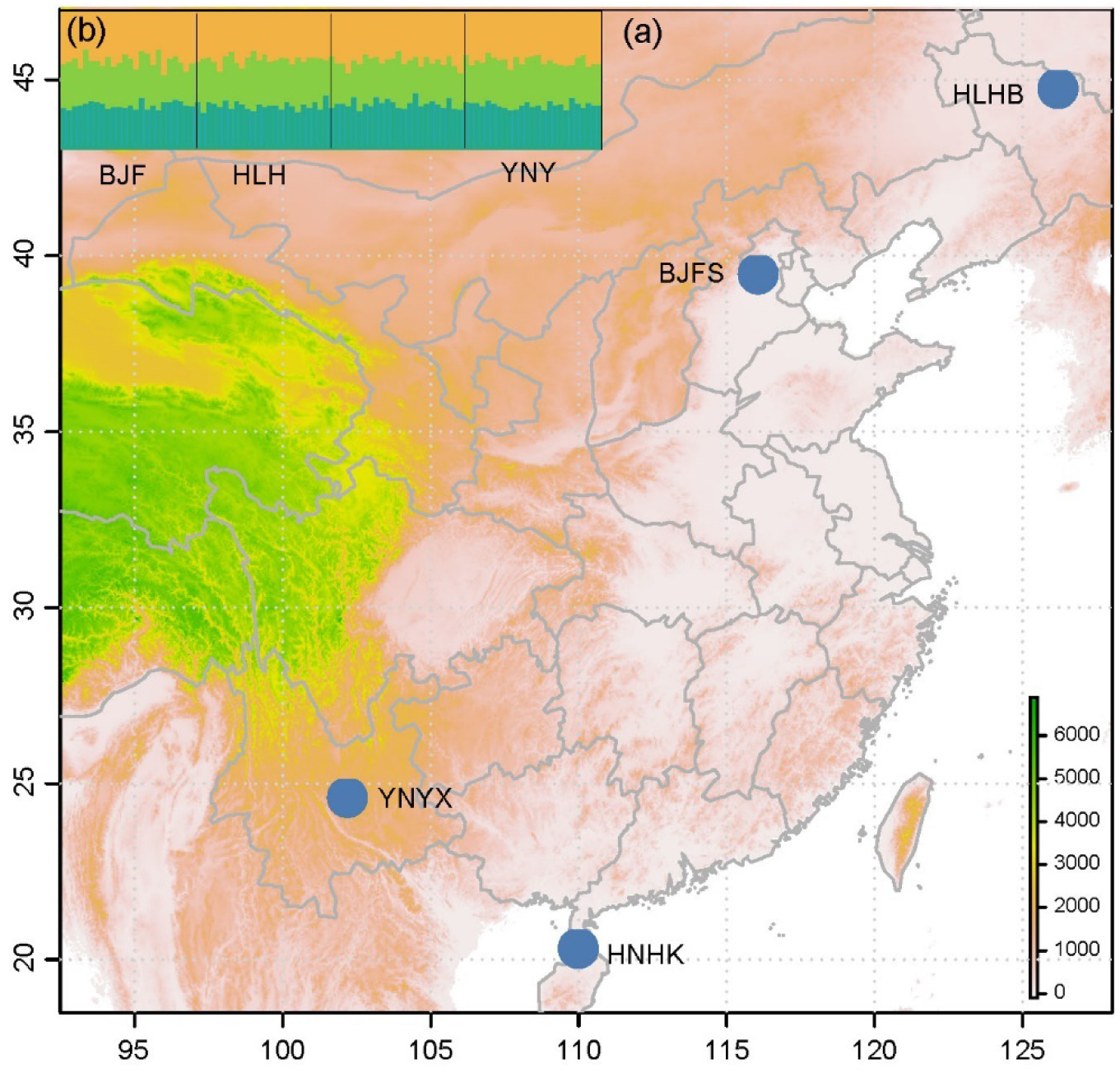
Collection sites of *Eupeodes corollae* (a) and population genetic structure analysis of four geographical populations using BAPS (a) and STRUCTURE (b). BAPS analysis showed that all population are clustered into one cluster (blue color in figure a). STRUCTURE analysis showed that the optimal delta K was three and all populations were composed of the three clusters. Codes for the population are shown in Table 1.

### Genome sequencing and assembly

The extracted genomic DNA from a laboratory-reared individual was used in constructing a high-throughput sequencing library with 500-bp insert size using the Illumina TruSeq DNA PCR-Free HT Library Prep Kit (Illumina, San Diego, CA, USA). The prepared library was sequenced on an Illumina Hiseq4000 Sequencer using the Hiseq Reagent Kit v3 (Illumina, San Diego, CA, USA) by Beijing BerryGenomics Co.,Ltd. The raw data were trimmed by removing the low quality reads using Trimmomatic 0.36 [24] and then the sequences were evaluated by FastQC v 0.11.5 [25]. IDBA was used to assemble the generated genomic sequences with K-mer from 20 to 140 [26].

### Genome-wide microsatellite survey and primer design

MSDB was used to search all potential microsatellite loci (repeat units of 2, 3, 4, 5, and 6 corresponding to the minimum number of repeats of 7, 5, 4, 4, and 4, respectively) from the assembled genomic sequences of *E. corollae* [27]. QDD was used to isolate microsatellites and design primers [28]. The outputs of primer pairs from QDD were further filtered by the following criteria [29, 30] : (i) the corresponding microsatellites were pure and specific; (ii) the design strategy of ‘A’ was used to avoid primer secondary structure and repeats; (iii) the minimum distance between the 3′ end of a primer and its target region should be longer than 10 bp; (iv) the annealing temperature for each primer pairs was set between 58 °C and 62 °C to avoid large differences among primers; (v) the estimated PCR product size of the primer pairs was from 100 to 350 bp.

### Polymorphic microsatellite isolation

After screening primers from the QDD program, a universal primer (CAGGACCAGGCTACCGTG) was added to the 5′ end of each selected forward primer to allow efficient combining with the fluorescent label [31]. Amplifications were performed using the GoTaq Green Master Mix (Promega, USA) in a final volume of 10 μl system with 0.5 μl of template DNA (5–20 ng/μl), 5 μl of Master Mix (Promega, Madison, WI, USA), 3.94 μl of ddH2O, 0.08 μl forward primer, 0.16 μl reverse primer and 0.32 μl universal primer labeled with fluorescence (FAM, HEX, and ROX sequencing dyes). The PCR protocol was set as: 5 min for 95 °C, 35 cycles of amplification with 95 °C for 30s, 56 °C for 40s, and 72 °C for 40s. Final extension was with 72 °C for 15 min. PCR products were analyzed on an ABI 3730xl DNA Analyzer (Applied Biosystems, USA) using the GeneScan 500 LIZ size standard (Applied Biosystems, USA). Genotyping was conducted by GENEMAPPER 4.0 (Applied Biosystems, USA). Those primer pairs with amplification efficiency lower than 75%, showing monomorphism in eight individuals, or producing more than two peaks (non-specific amplification) were discarded.

### Genetic diversity and population genetic structure analyses

GENEPOP version 4.0.11 [32] was used to test the likelihood of deviation from Hardy-Weinberg equilibrium (HWE) and the linkage disequilibrium (LD) at each microsatellite locus, the inbreeding coefficient (F_IS_) and pairwise population differentiation (F_ST_). Allele frequencies, expected heterozygosity (H_E_) and observed heterozygosity (H_O_) were calculated with the macros Microsatellite Tools [33].

Population genetic structure was analyzed by STRUCTURE version 2.3.4 [34]. The clustering test was replicated 30 times for each K value ranging from 1 to 5 with a burn-in of 100,000 iterations followed by 200,000 Markov Chain Monte Carlo iterations. The Delta (K) method was used to estimate optimal K values by submitting the STRUCTURE output to Structure Harvester Web 0.6.94 [35]. Visualization of the results was handled by CLUMPP version 1.1.2 [36] and DISTRUCT version 1.1 [37]. Additional, BAPS version 6.0 software (Bayesian analysis of population structure) was used to incorporate spatial information into clustering of individuals.

## Results and discussion

### Genomic sequences of E. corollae

A total of 51.53 Gb paired-end (PE) sequences (184,394,506 reads each with a length of 150 bp) was obtained and the genomic size of *E. corollae* was estimated to be 12315 Mb. Trimmed reads were assembled into 2563327 scaffolds with a total length of 1.15 Gb ranging from 100 bp to 437.63 KB, with an N50 of 1510 bp. These contigs were used for microsatellite discovery.

### Microsatellite characteristics of E. corollae

79,138 microsatellite loci were isolated from the randomly sequenced genome sequences of *E. corollae* with 5000 (6.32%) dinucleotide repeat (DNR) sites, 29221 (36.92%) trinucleotide repeat (TNR) sites, 30988 (39.16%) tetranucleotide repeat (TTNR) sites, 6635 (8.38%) pentanucleotide repeats (PNR) sites and 7294 (9.22%) hexanucleotide repeat (HNR) sites. The frequency of dinucleotide repeats in *E. corollae* is unusually low when compared with other insect species such as *Grapholita molesta* [30] (Lepidoptera), *Aphis glycines* (Hemiptera) [38] and *Obolodiplosis robiniae* (Diptera) [39], which shows the distribution of microsatellites to vary among species [40, 41].

### Development of variable microsatellite markers

The QDD program initially generated 18114 primer pairs; we selected those corresponding to tri-and tetra-nucleotide microsatellites for further filtering under criteria listed in the methods and obtained 40 primer pairs. These primer pairs were validated in eight individuals of *E. corollae*; six pairs with no polymorphism and ten pairs with low amplification efficiency (< 75%) were discarded. The remained 24 primer pairs that generated polymorphic genotypes were used for population-level examination.

Development of an appropriate set of markers is often the first step in population genetic and evolutionary studies. The recent development of genomic sequencing technology has made it relatively easy to isolate powerful microsatellites from large numbers of candidates at a genome-wide scale [42]. This method has been used in population structure analyses in many species, such as *Grapholita molesta* [30], *Frankliniella occidentalis* [43] and *Carposina sasakii* [29]. In our study, the 24 microsatellites developed are highly efficient in terms of amplification and polymorphism, enabling us to assess the population genetic structure of *E. corollae*.

### Population genetic diversity

A total of 96 individuals with 24 individuals from each of the four populations was used for the genetic diversity study. The number of alleles per locus for all individuals ranged from 5 to 13 with an average of 8.75, which showed the level of polymorphism of the selected loci. The observed (H_O_) and expected (H_E_) heterozygosity values ranged from 0.235 to 0.768 and from 0.333 to 0.785, respectively. Four loci (S01, S07, S24, S39) showed a significant gap between observed and expected values, while the inbreeding coefficient (F_IS_=(H_E_-H_O_)/H_E_) calculated by GENEPOP for these loci was relatively high.

Significant deviations from HWE after sequential Bonferroni correction [44] (P < 0.05) were detected in 9 of 24 loci (S01, S02, S07, S11, S14, S24, S25, S33&S39), and 3 of the 24 loci (S07, S24 & S39) deviated in all populations. None of the loci were in linkage disequilibrium (LD) in the four populations.

### Population genetic structure

Pairwise Fst analysis showed no significant differentiation between each pair of populations with F_ST_ values ranging from −0.007 to 0.001 (Table 4). BAPS analysis showed all populations clustered into one group (Figure 1a) while STRUCTURE analysis showed an optimal value of K=3. All populations were evenly spread across the three clusters, indicating a lack of genetic differentiation among populations (Figure 1b). This pattern of genetic structure is congruent with an estimated pairwise Fst values among populations. The geographically related pairs of populations had relatively small Fst value while the distantly related pairs of populations had relatively larger Fst values (Table 4).

**Table 4.** Pairwise Fst of 4 *Eupeodes corollae* populations based on 24 microsatellites.

| Population | BJFS | HLHB | HNHK | YNYX |
| --- | --- | --- | --- | --- |
| BJFS | — | 0.901 | 0.306 | 0.892 |
| HLHB | -0.004 | — | 0.838 | 0.973 |
| HNHK | 0.005 | -0.001 | — | 0.468 |
| YNYX | -0.004 | -0.007 | 0.001 | — |

The bottom triangle shows the pairwise Fst values, while the upper triangle shows the corresponding P values.

A lack of population differentiation is common in hoverflies. For example, a previous study on the hoverfly *Cheilosia naruska* from Finland showed that the species lacks differentiation at both the genetic and phenotypic levels [45]. Another study of two hoverfly species (*Episyrphus balteatus* and *Sphaerophoria scripta*) in Europe using 12 species-specific microsatellite markers also revealed a lack of genetic differentiation within species [18]. High levels of genetic diversity associated with a lack of structuring at a large spatial scale may indicate a high tolerance to environmental variability and a high migration rate [46]. Our study indicated that *E. corollae* in China may be highly mobile. The geographically related pattern of population structure may indicate that migration is restricted by geographical barriers. Our study provides preliminary insight into the biology and ecology of *E. corollae*. Further denser sampling is required to assess the population genetic structure of this species as well as other approaches to investigate its migration pattern.

Microsatellite markers are popular and powerful DNA markers because they are cost-effective and with a high diversity[41]. With the development of next-generation sequencing, genome-wide single nucleotide polymorphisms (SNPs) are becoming more powerful to screen genome-wide polymorphisms in a rapid and cost-effective manner [47]. Incorporating high-density SNPs in population genetic analysis may provide information on biology and ecology, such migration routes, of *E. corollae*, and help to understand adaptive evolution in this species [48].

## Conclusions

We developed 24 microsatellite markers in *E. corollae* at a genome-wide scale which provides genetic markers for population genetic analyses of this species. Our preliminary examination of four geographical populations of *E. corollae* across China suggested weak but geographically lined population differentiation. The results provide insight into migration of *E. corollae* in China.

## Acknowledgments

We thank Prof. Ary Hoffmann from The University of Melbourne for his revisions and suggestions on the manuscript, Xu-Bo Wang and Hua-Yan Chen for their help on the collection of specimens, Yan-Jie Lv for her help on the molecular works. This research was supported by the Applied Foundation of Science & Technology Department of Sichuan Province (2018YYJC0468), the National Natural Science Foundation (31472025), the Natural Science Foundation of Beijing Municipality (6162010), the Beijing Key Laboratory of Environmentally Friendly Pest Management on Northern Fruits (BZ0432), all of China.

## Author Contributions

Conceiving and design, Shu-Jun Wei and De-Qiang Pu; Data curation, L-Jun Cao, Ling Ma; Formal analysis, Meng-Jia Liu, Ling Ma and Li-Jun Cao; Methodology, Li-Jun Cao and Shu-Jun Wei; Resources, De-Qiang Pu, Ya-Jun Gong; Supervision, Shu-Jun Wei; Validation, Ya-Jun Gong and Li-Jun Cao; Writing – original random, Meng-Jia Liu, Ling Ma; Writing – review & editing, Meng-Jia Liu and Shu-Jun Wei.

